# Gene Symphony: Two-Layered Musification of Gene Expression Clusters with Functional Backgrounds

**DOI:** 10.1101/2025.06.13.659566

**Authors:** B Goutham, Manikandan Narayanan

**Author notes:** Contributing authors.

## Abstract

Musification, the process of converting data into music, has found applications in scientific research as a novel medium for data representation and communication. However, in biology, its application has largely been limited to DNA and protein sequences or structural data. Existing approaches that attempt to musify gene expression data often do so at the level of the entire gene set or its low-dimensional representations, lacking finer biological context. In our work, we bridge this gap by representing time-series gene expression data through clusters of genes that exhibit similar expression patterns across time points. We then embed contextual biological information onto these clusters to create layered sonifications: a foreground layer capturing expression dynamics across time points through pitch variations, and a background layer reflecting biologically meaningful context, such as Gene Ontology-derived semantic similarity, conveyed through smoothed audio modulated by intensity and echo effects. This musification approach offers an alternative way to analyze traditional visualizations like gene expression heatmaps and dendrograms, providing auditory insight into both temporal and functional trends. Preliminary user survey results indicate that 72.5% of participants were able to correctly interpret gene expression patterns from the foreground melody, and 88.9% perceived differences in background layers, highlighting the effectiveness of the layered musical representation in conveying biologically relevant information. This two-layered design is broadly applicable and can be extended to other forms of musification, allowing different types of information to be conveyed simultaneously through distinct musical components.

## 1 Introduction

### 1.1 Motivation

Sonification, the process of translating data into sound, is an innovative technique that has gained popularity across various scientific disciplines as an alternative method for data representation [1, 2]. By transforming quantitative information into auditory signals, sonification provides a new sensory channel through which individuals can perceive and analyze data. An extension of this concept, musification, specifically focuses on creating musical compositions from data, combining artistic expression with analytical utility. Traditionally, scientific research has relied heavily on visual data representations such as graphs and heatmaps to convey complex information. However, these visual representations can be limiting, particularly for those with visual impairments or those who find it challenging to interpret large volumes of information through traditional data presentation methods. Sonification, and by extension musification, bridges this gap by making data accessible through sound, offering a unique opportunity for alternative data analysis that can enrich our understanding.

### 1.2 Prior Works

Musification has found applications in biology. A review article by Braun et al. [3] discusses various studies that have explored its use in representing biological data, particularly focusing on DNA and protein datasets. Apart from genomic and proteomic data, sonification has also been used in biomedical research [4]. Gene expression data, especially time-series expression data, is an area in biology that has yet to be fully explored through musification. While a few studies have examined its use, there is still considerable room for further investigation and application in this domain (refer Literatur Review). Time-series gene expression data, typically represented through heatmaps, capture the dynamics of gene activity across various time points. However, heatmaps as visual representations are inherently limited as they primarily convey only the magnitude of expression changes and gene identities, without offering an intuitive way to integrate additional layers of biological meaning, such as functional similarity or semantic context.

### 1.3 Our Approach

Our method employs a two-layered musification strategy that transforms gene expression data into meaningful musical representations. Rather than musifying the entire gene set at once as done in earlier studies, we divide the gene set into clusters of genes that exhibit similar temporal expression patterns. By focusing on these clusters, our method captures the expression dynamics and provides an auditory representation of gene trends over time. This approach offers an alternative way to interpret heatmaps and dendrograms and makes the resulting music accessible and intuitive.

In addition to expression patterns, we introduce a second dimension of biological context to the music by incorporating functional enrichment analysis using Gene Ontology (GO) terms. These GO terms characterize the biological functions and processes associated with each gene cluster. To embed this biological context, we designed a layered music generation framework. The foreground layer encodes gene expression dynamics using pitch — where higher expression values produce higher notes — while the background layer encodes the semantic similarity of GO terms associated with the cluster. The background music varies in intensity and echo strength depending on the semantic cohesion of the cluster, allowing listeners to audibly sense the degree of biological relatedness among the genes.

To evaluate the effectiveness of our two-layered musification approach, we conducted a survey involving 18 participants — 13 with a background in computational biology and 5 without prior domain knowledge. The survey tested multiple aspects of interpretability across different musification strategies, including our Chords and Mean methods. Participants first matched audio clips to expression plots, achieving a 72.5% accuracy across three samples, indicating that expression trends were meaningfully conveyed. In the second task, where listeners identified the correct gene cluster from individual gene audio, an average accuracy of 63.9% across both strategies demonstrated the ability to perceive expression dynamics. In a final perception task, 88.9% of participants correctly detected the presence of echo in the background layer, confirming that semantic information — driven by Gene Ontology-based enrichment — was effectively communicated through sound.

This layered musification framework not only allows scientists to explore gene expression from a new perspective but also introduces a multidimensional auditory representation that blends expression behavior with semantic context. The concept of using distinct musical layers to represent different kinds of information can be extended to other types of musification tasks across domains — offering a generalizable approach where one layer encodes primary data features, and another conveys contextual or meta-information.

The beneficiaries include not only scientists and researchers who can use this method for biological analysis, but also individuals who may find the auditory representation of data more accessible, such as those who are visually impaired or color-blind. By providing an alternative form of data analysis through music, we open up new avenues for understanding biological phenomena. Furthermore, the aesthetic nature of the music encourages engagement with the data even from individuals with no prior interest in science.

## 2 Literature Review

### 2.1 Related Works

#### 2.1.1 Musification of DNA & Protein Sequences

Sonifying genomic and proteomic sequences has intrigued researchers for decades. One of the earliest notable efforts in this field was John Dunn’s presentation, “Inflections: Musical Interpretations of DNA Data,” at a University of Hawaii conference in the late 1980s [5, 6]. Since then, numerous studies have explored the interpretation of sequence data through audio representation [7–9]. One of the recent works by Martin et al. [10] presents sonification techniques to analyse protein sequence data. They implemented five sonification algorithms designed to map protein data characteristics into sound, each with a unique approach for single protein sequences or Multiple Sequence Alignments. The first three algorithms focus on sonifying individual protein sequences based on hydrophobicity, where Algorithm I maps amino acids to specific MIDI (Musical Instrument Digital Interface) pitches using the Goldman-Engelman-Steitz hydrophobicity scale, allowing listeners to recognize patterns through pitch variation. Algorithm II further simplifies this by grouping amino acids into four hydrophobicity-based clusters and Algorithm III combines both approaches by applying the detailed mapping of Algorithm I while using separate instruments for each group from Algorithm II. The fourth and fifth algorithm focus on sonifying multiple sequence alignments (MSAs) to reflect conservation patterns across sequences. Algorithm IV uses Shannon entropy to assign pitch based on the level of conservation at each position in the MSA, with higher pitches indicating less conservation and Algorithm V generates polyphonic output by mapping the presence of amino acids across MSA positions to pitch and adjusting volume based on consensus, highlighting sequence consistency within the alignment

#### 2.1.2 Musification of Expression Data

In contrast to the sonification of sequence data, relatively very few studies have focused on the sonification of gene expression data.

##### Static Gene Expression Data

In prior research, Staege [11] introduced GEMusicA algorithm, a novel method to represent gene expression data through musical compositions. GEMusicA translates static gene expression levels, using microarray data from four neuroblastoma-derived cell lines, into musical elements like pitch (the frequency or the note) and tone duration (the time each note is played), providing an intuitive way to visualize and analyze gene expression patterns. Each musical note is calculated by dividing the intensity of a microarray probe set (gene) by the maximum intensity within the set, then scaling it based on the instrument and rounding to the nearest whole number. The probe sets with high variability (top 10% in their study) are selected to highlight the differences more clearly. Tones are scaled based on frequency (e.g., higher frequencies represent higher signal intensities), and duration is adjusted to reflect the information content of each gene, with genes showing high variance rendered as longer notes. To facilitate better comparison between samples and enhance interpretability for non-musicians, the frequencies are re-normalized using a “reference melody”. The reference melody, usually well-known piano pieces, is then used to tweak the earlier calculated median frequencies of a probe set across all samples.

While this approach offers a creative means to interpret biological data, with applications in fields such as stem cell biology and cancer research [12], it encounters technical issues related to calculation precision and limited musical parameters, which may present a barrier for researchers unfamiliar with music. Furthermore, this work does not emphasize robust clustering techniques for probe sets (genes) to reduce the dataset before mapping it to the musical scale. By filtering the original dataset of 22,283 probe sets down to 192 through low-variance thresholds [13], this method risks data loss, potentially undermining the goal of representing the full range of gene expression data in musical form.

##### Time-Series Gene Expression Data

Another work by Alterovitz and Yuditskaya [14] utilizes sonification techniques to map gene expression and protein abundance patterns across time to musical frequencies. This is done through principal component analysis (PCA), which reduces the dimensionality of gene expression data and identifies the key components representing the most significant variations in gene activity. These principal components are then mapped to musical intervals, using Pythagorean tuning principles to ensure that control data produces harmonious music, while experimental or abnormal data creates dissonant sounds. The method is applied to a colon cancer dataset, demonstrating that normal gene expression patterns produce harmonious musical sequences, while cancer samples generate inharmonious sounds, reflecting underlying changes in gene dynamics. This comparative sonification approach provides an alternative, auditory method for real-time monitoring of gene expression and protein abundance, which could be useful for applications like personalized medicine and patient health monitoring. By incorporating the use of sound for real-time tracking of cellular protein levels and gene expression, the method enables physicians and researchers to identify irregularities without depending on visual data, thereby reducing cognitive strain and enhancing accessibility.

### 2.2 Background

#### 2.2.1 Over Representation Analysis (ORA)

Over Representation Analysis (ORA) is a widely used statistical method in functional enrichment studies to identify whether specific biological categories, such as pathways or Gene Ontology (GO) terms, are disproportionately represented within a set of genes of interest compared to a background set. The core idea involves constructing a contingency table to test whether the number of genes from a particular category observed in the input list is significantly higher than expected by chance. This is typically assessed using statistical tests like the hypergeometric test or Fisher’s exact test, which compare the observed count of genes associated with a category in the target list to the expected count based on the background. Several tools have been developed to implement ORA, including DAVID [15], g:Profiler [16], and Enrichr [17]. In our project, WebGestalt (WEB-based Gene SeT AnaLysis Toolkit) is used, which provides an intuitive interface, customizable background settings, and support for various organisms and enrichment databases [18]. WebGestalt also integrates advanced features such as weighted set cover filtering to reduce redundancy among enriched terms, improving interpretability of results. This makes it particularly suitable for large-scale genomic analyses and for drawing biologically meaningful insights from gene expression data.

Recent studies have effectively applied ORA to identify biologically meaningful patterns in colorectal cancer (CRC). Zhao et al. [19] analyzed gene expression data from early-onset CRC patients and used functional enrichment tools, including DAVID, to reveal that differentially expressed genes were significantly associated with muscle contraction and vascular smooth muscle contraction pathways. In a related study, Huang and Cui [20] employed WebGestalt to perform GO and KEGG enrichment analysis on genes differentially expressed between colorectal cancer and adenoma samples, identifying key hub genes involved in immune response and chemokine signaling pathways. These applications of ORA demonstrate its effectiveness in identifying biological processes, relevant pathways, and molecular signatures, thereby aiding in the discovery of underlying mechanisms driving gene expression changes.

#### 2.2.2 Musical Instrument Digital Interface (MIDI)

The Musical Instrument Digital Interface (MIDI) is a communication standard that enables electronic instruments, computers, and other equipment to interact and exchange musical information. Rather than recording audio directly, MIDI captures the instructions for music, including notes, timing, and velocity, which can be interpreted and reproduced by compatible devices. Each musical note in MIDI is assigned a specific number, known as the MIDI note number, which corresponds to a particular pitch or frequency. For example, middle C on a piano is represented by MIDI note number 60, with each successive note assigned a unique MIDI number and its own frequency. In our study, the piano is used as the primary instrument for mapping MIDI data due to its tonal clarity and broad frequency range, making it an ideal choice for producing precise and distinguishable notes that facilitate accurate auditory representation of data-driven sounds.

## 3 Problem Statement

Sonification has been applied extensively throughout science to answer various research questions, but its application in biology is mostly limited to sequence and structure data. Although previous works by Staege [11], and Alterovitz and Yuditskaya [14] on gene expression data yielded insights—Staege [11] through static representations with GEMusicA for specific cell lines, and Alterovitz and Yuditskaya [14] via realtime sonification of time-series data with PCA for pattern detection—both approaches primarily focus on simplifying expression values across the *entire* gene set into a *single* musical representation using dimensionality reduction techniques, which may lead to data loss and overlook localized expression dynamics. Moreover, none of the studies integrate the biological information of the genes considered in their analysis.

We bridge this gap by introducing a sonification mapping algorithm and music generation techniques that represent and analyze time-series gene expression through *‘clusters’*, offering an alternative approach to interpreting gene expression heatmaps and dendrograms. By focusing on clusters, our method captures local expression dynamics, providing auditory insights into gene behavior over time. In addition to this, we incorporate functional enrichment analysis using Gene Ontology (GO) terms to characterize each gene cluster with biologically meaningful interpretations. The results of this analysis are then used to generate background music based on the semantic similarity of GO terms in each cluster. Specifically, our aim is:

*To represent an expression vector heatmap using musical outputs as a novel way of analyzing time-series gene expression clusters with functional enrichment results.*

To achieve the proposed goal, we break down the problem statement into an Input-Output pair.

**Input:** A time-series gene expression matrix **X** ∈ ℝ*^G^*^×*T*^, where *G* is the number of genes and *T* is the number of time points.

**Output:** Multiple clusters {𝒢_1_, 𝒢_2_*, …,* 𝒢*_C_* } of total *C*, where 𝒢*_i_* ⊆ 𝒢, and corresponding musical audio representations for each cluster *i*, containing information about expression dynamics over time as foreground music using one of the three methods:

- **Chords:** 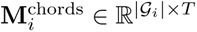 ; where notes at each time point *t* ∈ [1*, T* ] are played simultaneously with fixed duration Δ*t*.
- **Mean:** 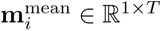, where

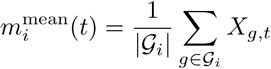 played sequentially with fixed duration Δ*t*.
- **Delayed:** 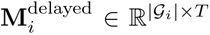 ; where notes at each time point *t* are sorted and played with intra-time delay *δ* with note onset of *t* + *j* · *δ*, *j* ∈ [0, |𝒢*_i_*| − 1].

and background music modulated using intensity variations and echoes using biological context, such as the semantic enrichment score *s_i_* ∈ [0, 1].

The total number of clusters *C*, can be chosen based on the user’s preference by changing the appropriate parameters during hierarchical clustering (refer Methodology).

We aim to evaluate the audio outputs generated through our methodologies to determine whether this novel approach can serve as an alternative representation and analysis of gene expression matrices. By transforming expression data into sound, we offer an accessible format that could be particularly valuable for visually impaired and color-blind individuals, enabling them to perceive variations in gene expression traditionally visualized in heatmaps. Additionally, this method brings an aesthetic dimension to biological data, allowing listeners to engage with gene expression patterns in a new sensory modality that combines scientific rigor with an artistic experience. Through this analysis, we hope to explore the potential of auditory representations to communicate biological information in a way that is both informative and inclusive.

## 4 Methodology

### 4.1 Pre-processing: Clustering

We applied hierarchical clustering using the Ward linkage method [21] to group genes based on similar temporal expression patterns. First, the dataset, which contained gene expression levels across multiple time points, is standardized using a z-score transformation to ensure comparability across genes. The hierarchical clustering process is then executed on the standardized data, with the Ward method selected for its ability to minimize intra-cluster variance. To define the clusters, we set a distance threshold in the hierarchical dendrogram, aiming for clusters of approximately 10-30 genes. This threshold is adjusted iteratively to achieve the desired cluster size, facilitating more interpretable groups while ensuring manageable cluster volumes.

To evaluate the internal coherence of the clusters, we calculate the intra-cluster similarity scores based on pairwise correlations among gene expression profiles within each cluster. These similarity scores, where higher values indicate stronger similarity, are distinct from the semantic similarity scores used later in the analysis; they are derived solely from expression values and not from gene functions. This metric is used to identify and rank clusters based on expression coherence, allowing us to select the top five to six clusters with the highest internal similarity for detailed analysis. For these selected clusters, we generate individual plots displaying gene expression patterns across time, with each gene labeled by its identifier to aid in observing distinct expression trajectories. This clustering methodology enables the identification of groups of genes with similar temporal expression patterns.

### 4.2 Foreground Layer

#### 4.2.1 Mapping To Musical Space

We use the Absolute Elevation Mapping strategy defined by Gao et al. [22] as our mapping function for the foreground music. In this strategy, the mapping of gene expression values to MIDI notes involves normalizing the expression values to fit within the instrument’s (in our case, the piano’s) MIDI note range. The minimum and maximum expression values across all clusters (*V*_min_ and *V*_max_) are identified to scale the values accordingly. Using these limits, the equations (1) and (2) transforms each expression value into a MIDI note by mapping it proportionally between MIDI MIN (21, representing A0 note on the piano) and MIDI MAX (108, representing C8 note on the piano), with *α* and *β* calculated to define this MIDI range. For each gene’s expression value *V*, the mapped note is calculated, and if this value is not an integer, rounding is applied based on its fractional component. If the fractional part is less than 0.5, it is rounded down with floor, while values with fractional parts of 0.5 or more are rounded up with ceil.

Mathematically, it can be formulated as:

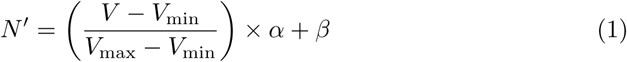

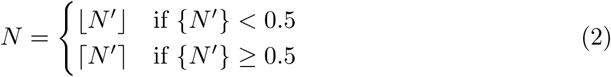

where:

- *N* is the MIDI note of the gene
- *V* is the expression value of the gene
- *α* is set arbitrarily based on the instrument (= 87 for the piano)
- *β* is a bias parameter which ensures notes are in the MIDI range (= 21 for the piano)
- *V*_min_ is the minimum expression value across all clusters
- *V*_max_ is the maximum expression value across all clusters

This approach ensures that expression values are converted to whole-number MIDI pitches while preserving the variations in the data. The resulting MIDI notes are stored in a new dataset for each cluster and used to create a unique MIDI file that represents the cluster’s gene expression dynamics as music.

#### 4.2.2 Generating Foreground Music

To generate foreground music from gene expression data, we’ve implemented three unique approaches: “Chords”, “Mean”, and “Delayed”. Each approach takes clusters of gene expression values, maps them onto MIDI notes scaled within the piano’s range (notes 21 to 108), and assigns these notes to represent specific pitches.

##### Chords

In the “Chords” method, all gene expressions within a cluster at each time point are represented simultaneously, forming a chord. Let 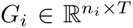 be the expression matrix for the *i^th^* cluster, where *n_i_* is the number of genes and *T* the number of time points. For each *t* ∈ {1, 2*, …, T* }, all *n_i_* expression values *G_i_*[:*, t*] are mapped to corresponding MIDI notes 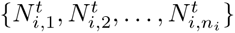 and played simultaneously with a fixed duration Δ*t*. Each time point corresponds to a new chord, with each gene mapped to a note in the chord. This approach allows all genes in the cluster to be “heard” together, capturing collective expression patterns as they evolve over time.

##### Mean

The “Mean” method calculates the mean gene expression value for each time point in each cluster. For 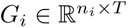, the mean vector is 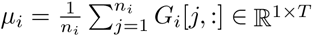. Each *µ_i_*[*t*] is mapped to a single note 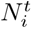. Instead of individual gene notes, a single representative note is generated for each time point based on this mean expression level and played with fixed duration Δ*t*. This creates a more averaged sound that reflects the overall trend of gene expressions rather than individual fluctuations.

##### Delayed

The “Delayed” method introduces temporal staggering to simulate a cascading auditory effect. For each time point *t* ∈ {1, 2*, …, T* } in a given cluster matrix 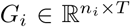, the gene expression values *G_i_*[:*, t*] are sorted in ascending order and mapped to MIDI notes. Instead of playing all notes simultaneously, each note is assigned an onset time offset by a delay *δ*. Specifically, the *j*^th^ note (from the sorted list) is played at time *t* + *j* · *δ*, where *j* ∈ {0, 1*, …, n_i_* − 1}. After each time point, the global MIDI time advances by a fixed increment. This method provides a unique “arpeggio-like” pattern, where each gene’s expression level is heard in sequence with a slight delay, allowing individual notes to stand out within the context of each time point’s chord.

These methods allow each gene’s expression to be translated into sound, creating distinct auditory patterns for each cluster. While all methods use the same data-to-note mapping technique, they differ in how notes are arranged and timed, offering varied musical interpretations of the data.

### 4.3 Background Layer

#### 4.3.1 Functional Enrichment using WebGestalt

To functionally interpret each gene cluster, we performed Over-Representation Analysis (ORA) using the WebGestaltR package [18]. Each cluster was analyzed independently against a common reference set consisting of 15271 protein-coding genes from the murine dataset (see Data Section). Only Gene Ontology (GO) terms with a false discovery rate (FDR) below 0.05 were considered statistically significant. The FDR was controlled using the Benjamini-Hochberg (BH) procedure [23], which adjusts raw *p*-values to account for multiple hypothesis testing. The BH method ensures that the expected proportion of false positives among the reported significant results remains below the 5% threshold.

Among the statistically significant GO terms, the top three were selected based on the *enrichment ratio*, defined as: Enrichment Ratio =

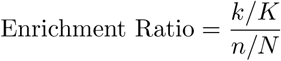

where:

- *k* is the number of genes in the input cluster that fall under a given GO term,
- *K* is the total number of genes in the input cluster,
- *n* is the number of genes in the background that fall under that GO term,
- *N* is the total number of genes in the background reference.

A higher enrichment ratio indicates a stronger overrepresentation of a GO term in the input gene cluster.

#### 4.3.2 Semantic Score using Wang’s Method

To evaluate the functional coherence of enriched GO terms obtained from each gene cluster, we calculated a semantic similarity score. This score quantifies how functionally related the top enriched Gene Ontology (GO) terms are, using the Biological Process (BP) functional database.

To quantify the semantic similarity among the top three enriched GO terms for each cluster, we used the GOSemSim R package [24]. Specifically, we used Wang’s method [25], which relies on the structure of the GO Directed Acyclic Graph (DAG) to compute similarity.

In Wang’s method, each GO term *t* is represented as a DAG of itself and its ancestor terms in the ontology. The contribution of each ancestor term *a* to *t* is calculated as:

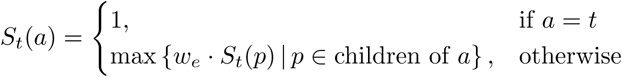

where *w_e_* is a semantic contribution factor assigned to the type of relationship between GO terms (e.g., ‘is-a’ or ‘part-of’ in Figure 3).

**Fig. 1.**
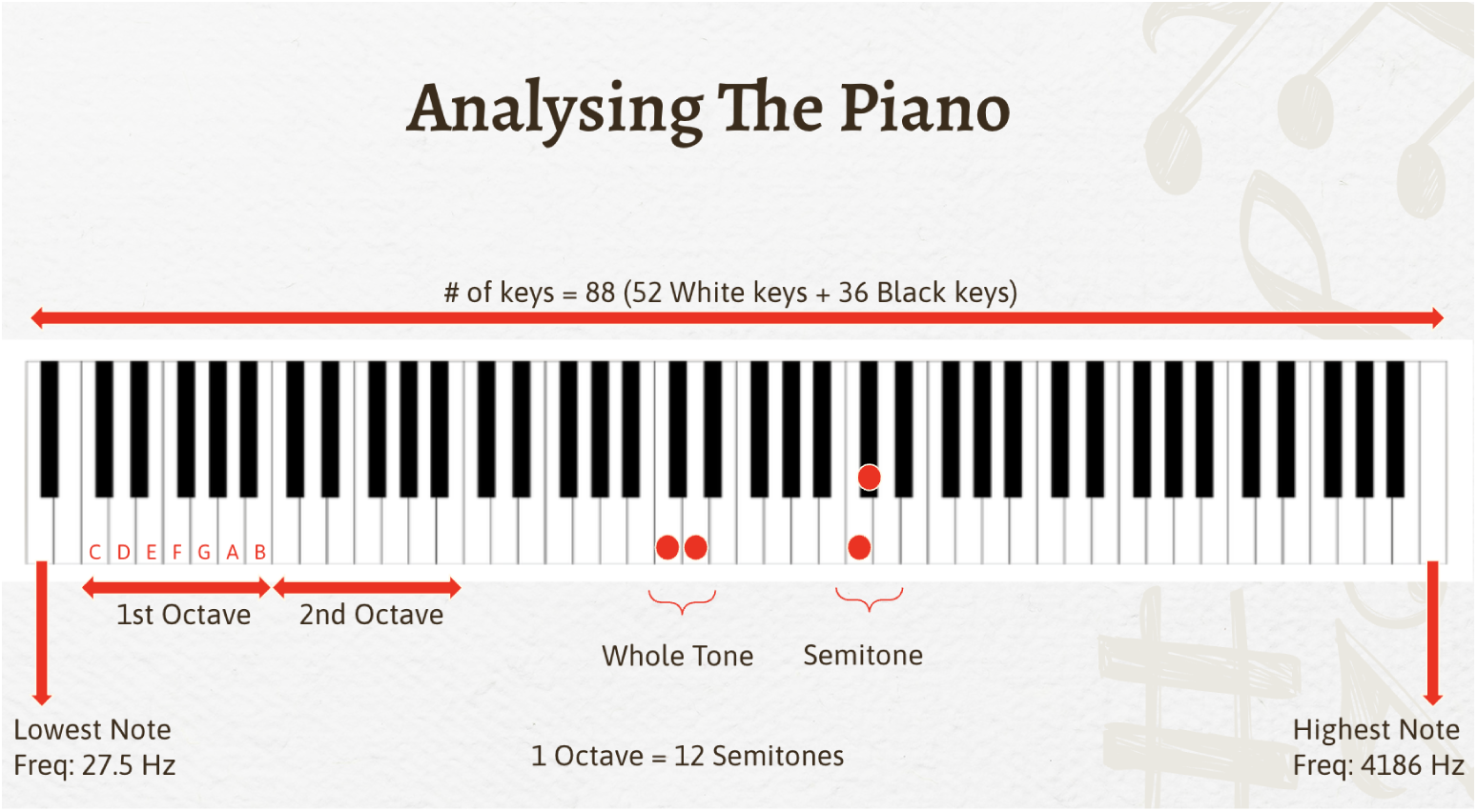
The piano contains a total of 88 keys. It is divided into multiple octaves with keys C, D, E, F, G, A and B. The frequency corresponding to the lowest note is 27.5 Hz, while the highest note corresponds to 4186 Hz.

**Fig. 2.**
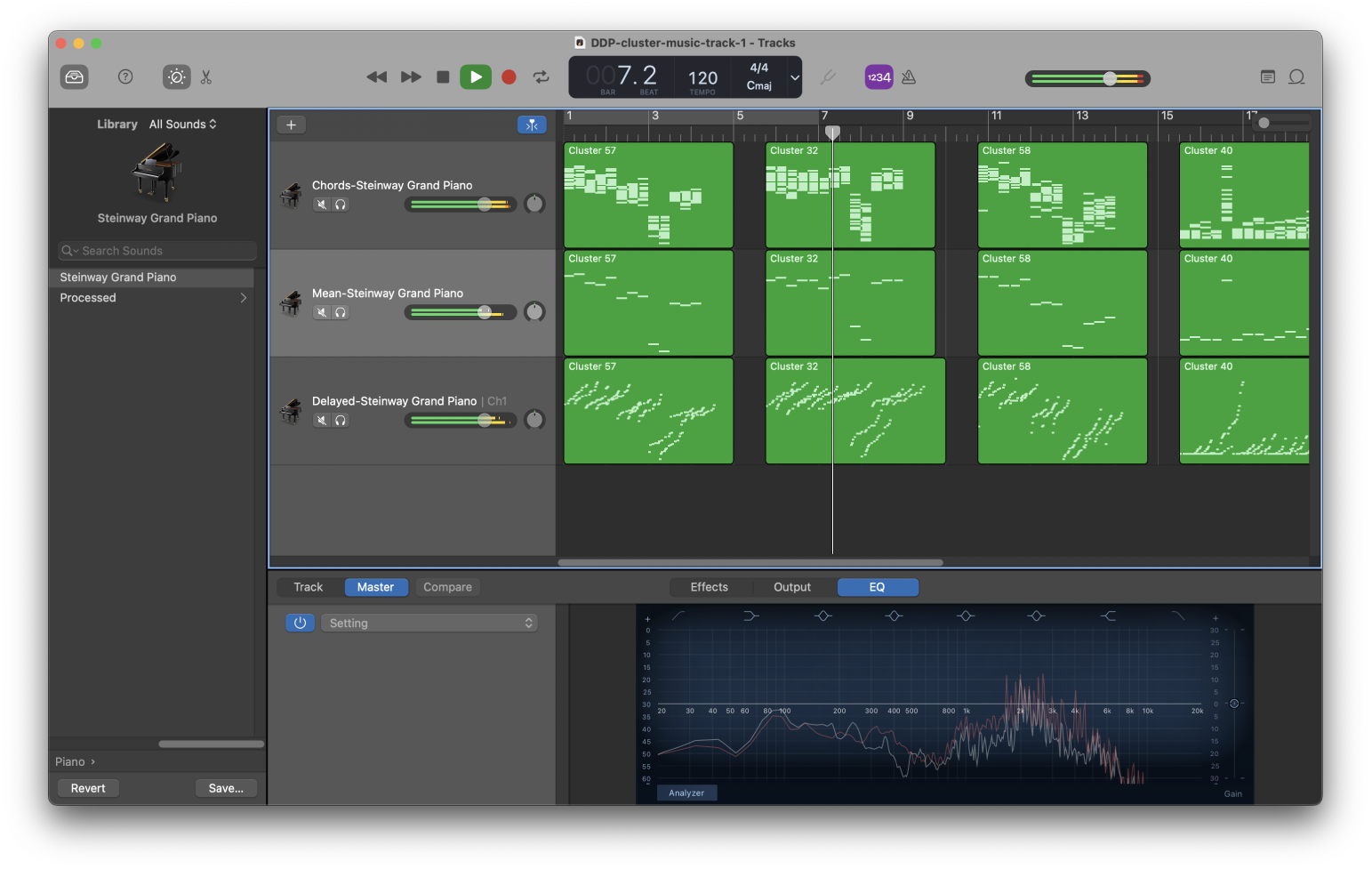
Utilizing the GarageBand software to visualize the three methods used to generate foreground music for each cluster. The top row represents the Chords method, where each time point has |𝒢*_i_*| notes, with |𝒢*_i_*| being the number of genes in the cluster 𝒢*_i_*. The middle row represents the Mean method, where one note corresponding to the average of expression values of genes is played at each time point. The bottom row represents the Delayed method, where notes are played with a small delay between the notes of the Chords method at each time point in ascending order.

**Fig. 3.**
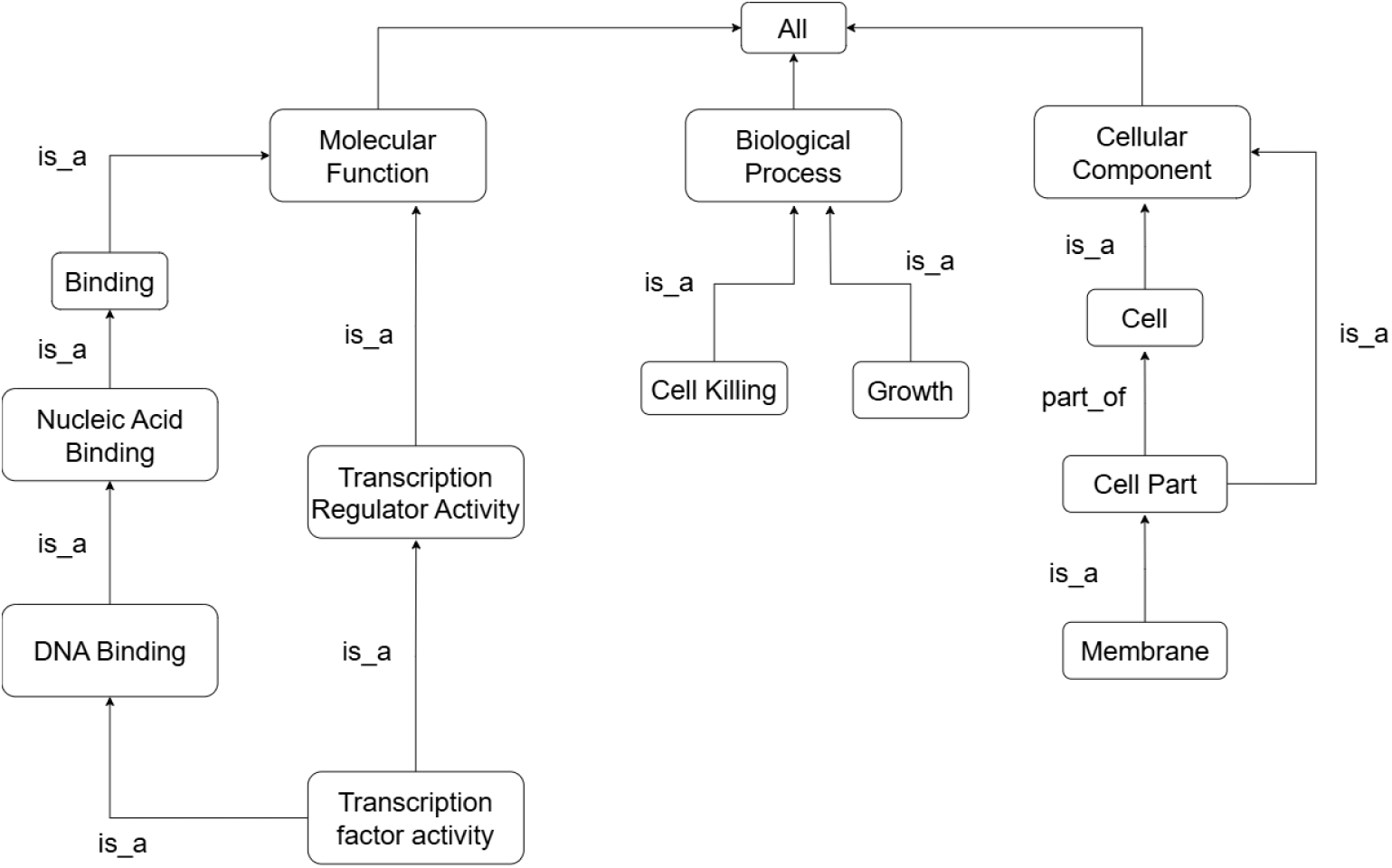
An illustrative Directed Acyclic Graph (DAG) depicting the hierarchical structure of Gene Ontology (GO) terms. The graph shows relationships among terms from the three main GO domains—Molecular Function, Biological Process, and Cellular Component—connected through ‘is-a’ or ‘part-of’ relationships. This structure forms the basis for semantic similarity calculations, such as Wang’s method, which quantify functional relatedness between GO terms based on their positions and connections within the GO hierarchy. Image redrawn from Almasoud et al. [26]

The semantic value of a GO term *t* is:

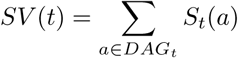

Given two GO terms *t*_1_ and *t*_2_, the Wang similarity is computed as:

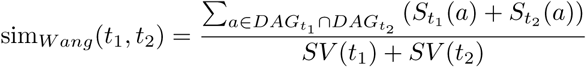

For each cluster, the three most enriched GO terms (based on enrichment ratio) were selected. All pairwise combinations (i.e., ^3^ = 3 pairs) of these GO terms were used to compute the similarity scores. The final semantic score for the cluster is defined as the mean of these pairwise similarity scores:

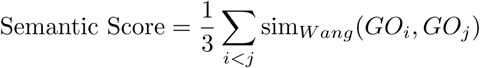

This average score captures how functionally cohesive the top enriched biological processes are within each cluster. Higher scores indicate more semantically consistent functional annotations, suggesting more biologically meaningful groupings.

#### 4.3.3 Generating Background Music

We introduce background music that is contextually informed by functional enrichment analysis. Each gene cluster is subjected to semantic analysis based on its enriched biological functions, resulting in a *semantic score* that influences the characteristics of the background audio. This score serves as a proxy for the biological “intensity” or “importance” of each cluster and modulates the smoothness and echo in the background sound.

##### Background Smoothing via Frequency Interpolation

To generate a continuous and fluid background audio track, we employed linear interpolation over the discrete sequence of frequencies derived from mapped MIDI notes of foreground music. These MIDI notes were first converted into frequencies using the standard formula:

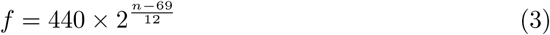

where *n* denotes the MIDI note number, and *f* is the corresponding frequency in Hertz. Given a time series of frequencies *f*_0_*, f*_1_*, …, f_T_*, we interpolate them over a denser time grid of length *L* to simulate a smooth pitch transition. Let *L* = *T* · *D* · *f_s_*, where *D* is the duration per note (in seconds), and *f_s_* = 44100 Hz is the sample rate. We define:

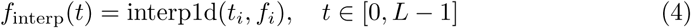

where *f*_interp_(*t*) is obtained via linear interpolation of the original frequencies *f_i_* at discrete indices *t_i_*. The interpolated frequency sequence is then used to generate the waveform using a cumulative phase approach:

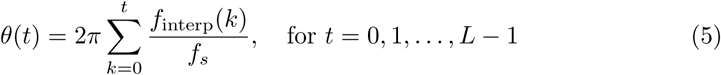

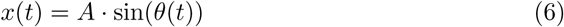

where the amplitude *A* varies linearly with the semantic score in the lower range and saturates beyond a threshold.. This results in a smooth sinusoidal signal whose frequency changes continuously over time, producing a fluid background ambiance.

##### Echo Addition for Strong Coherence

To add more significance to the background audio for high semantic scores, we implemented a multi-repeat echo effect. Echoes were simulated by repeatedly attenuating and delaying the original signal. Let *x*(*t*) denote the original audio signal, and let *d* be the fixed delay in samples. We define the echo-enhanced signal *y*(*t*) as:

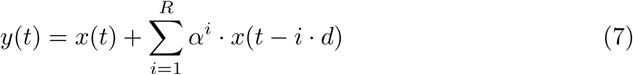

where:

- *R* is the number of echo repeats,
- *α* is the echo attenuation factor (0 *< α <* 1),
- *d* is the delay in samples, given by *d* = delay_sec_ × *f_s_*,
- *f_s_* is the sample rate.

In our implementation, we used *R* = 4 repetitions and a fixed delay of 0.3 seconds between each. The echo strength *α* is scaled non-linearly using a quadratic function of the semantic score, such that higher semantic scores yield stronger and more noticeable echoes. For each repetition, the echo is attenuated by a factor of *α^i^*, where *i* is the repetition index. This recursive layering of delayed and attenuated signals creates a trail of decaying echoes, enriching the background audio to reflect greater semantic relevance.

The final audio for each cluster is a combination of foreground music—generated using one of the previously described methods (Chords, Mean, Delayed)—and a continuous background audio following the frequency pitches of the foreground audio across time points. The background audio is further added with an echo for higher semantic scores. The semantic score obtained from enrichment analysis modulates the expressiveness of the background layer, adding a functional annotation component to the auditory experience.

## 5 Results

### 5.1 Data Used

The dataset used in this project was obtained from the study *TiSA: TimeSeriesAnalysis* by Lefol et al. [27], which provides a comprehensive pipeline for the analysis of longitudinal transcriptomics data. The data comprises mRNA-seq profiles from murine primary B-cells stimulated with LPS, IL-4, and TGF-*β*1, collected at ten time points post-stimulation: 0 h, 15 min, 30 min, 1 h, 2 h, 3 h, 6 h, 12 h, 24 h, and 48 h, each with a matched unstimulated control. They performed sequencing read alignment to the mouse reference genome using STAR [28], and quantified gene-level counts using htseq-count [29].

For our analysis, only the target expression matrix from the dataset was used. This matrix initially contained 41115 murine gene entries. To ensure suitability for downstream enrichment analysis, the dataset was filtered to retain only protein-coding genes, resulting in a set of 18027 genes. Additionally, any genes exhibiting zero expression across all time points were removed. The final set contained 15271 murine genes.

We present the results obtained using our methodologies on the murine time-series gene expression vector. We begin with the identification of similarly expressed gene clusters using hierarchical clustering. This is followed by functional enrichment analysis using WebGestalt to uncover the underlying biological relevance of each cluster. Next, we describe the music generated from these expression dynamics using multiple musification strategies. Mel spectrograms are provided to visualize the structure of the audio outputs. Lastly, we report findings from a structured user survey designed to assess the clarity, interpretability, and perceived differences in the generated musical representations.

### 5.2 Clusters and Enrichment Results for Murine Data

Figure 4 shows the expression pattern for a cluster after performing the Ward’s linkage hierarchical clustering on the murine time series expression data containing 15271 genes and 10 different time points. Each colored line represents the temporal expression profile of an individual gene within the cluster. The prominent black line signifies the mean expression level across all genes at each time point, while the shaded grey region denotes the two standard deviation confidence interval, capturing the variability within the cluster. Figure 5 shows similar plots for the six other gene clusters produced.

**Fig. 4.**
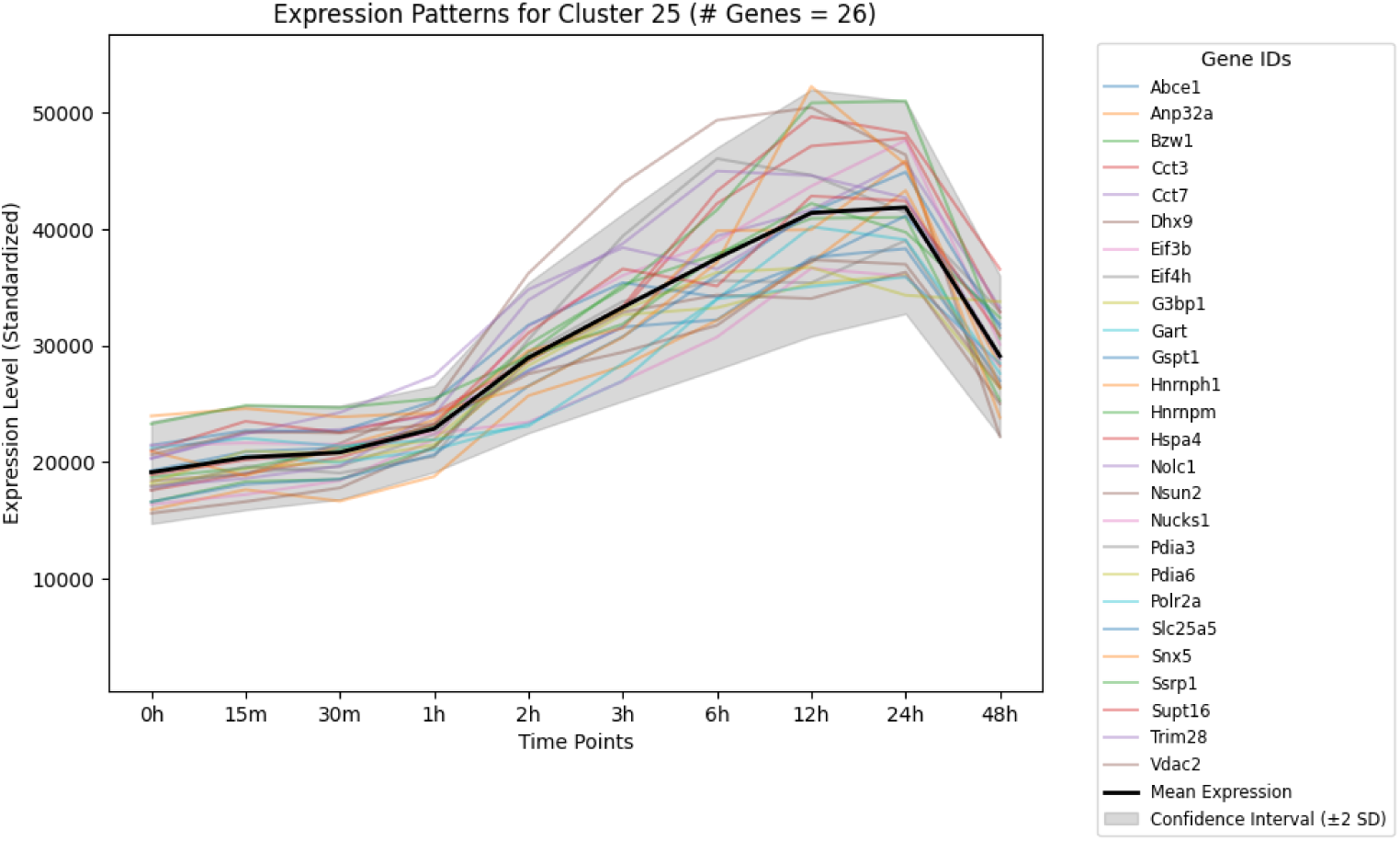
Expression Patterns for a Cluster (label 25) containing |𝒢*_i_*| = 26 genes obtained after performing hierarchical clustering on the murine expression vector containing *T* = 10 time points.

**Fig. 5.**
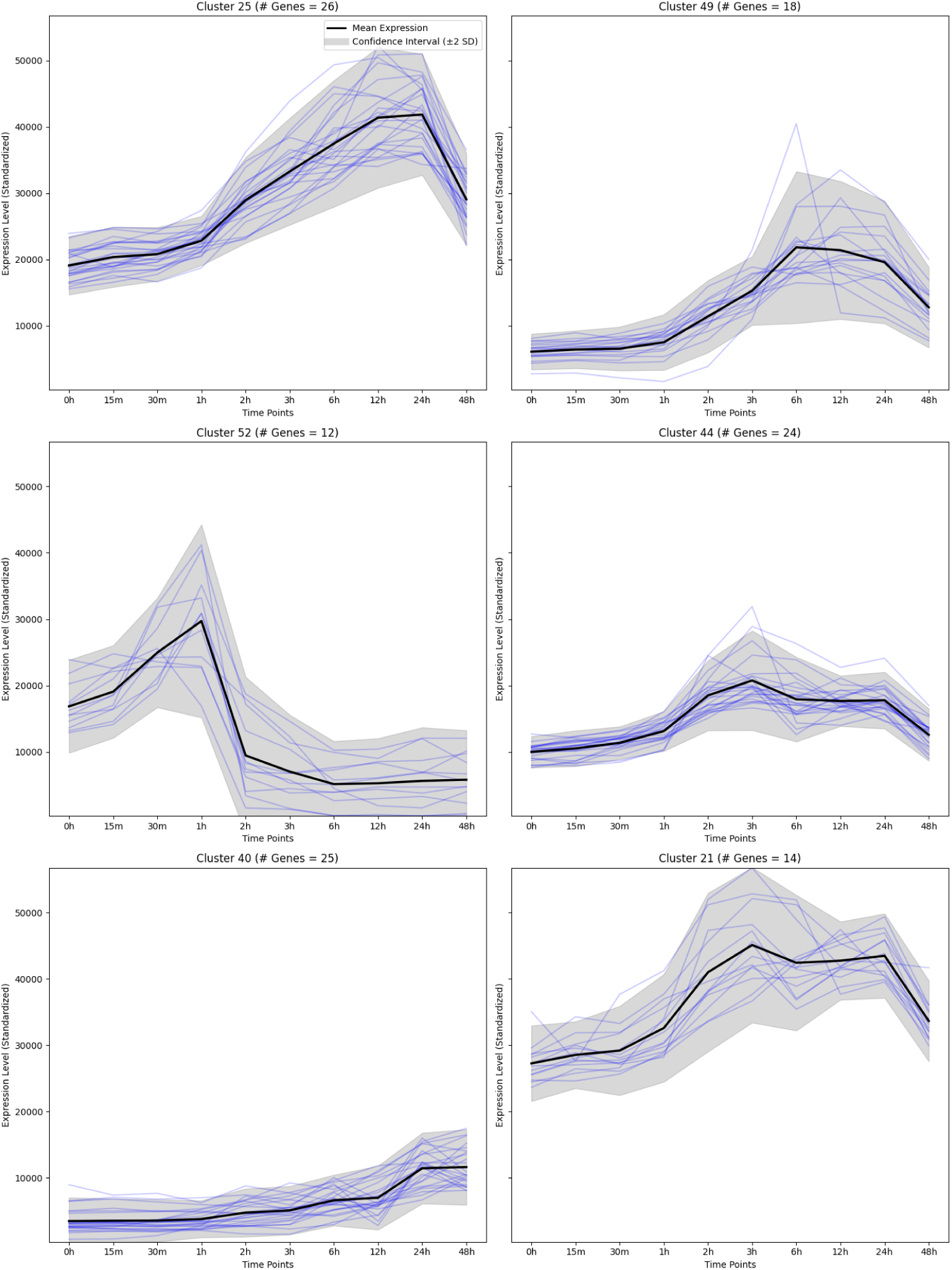
Expression patterns for multiple clusters with total number of genes, |𝒢*_i_*|, between 10-30 from the murine expression vector. These clusters are further used for enrichment analysis.

**Fig. 6.**
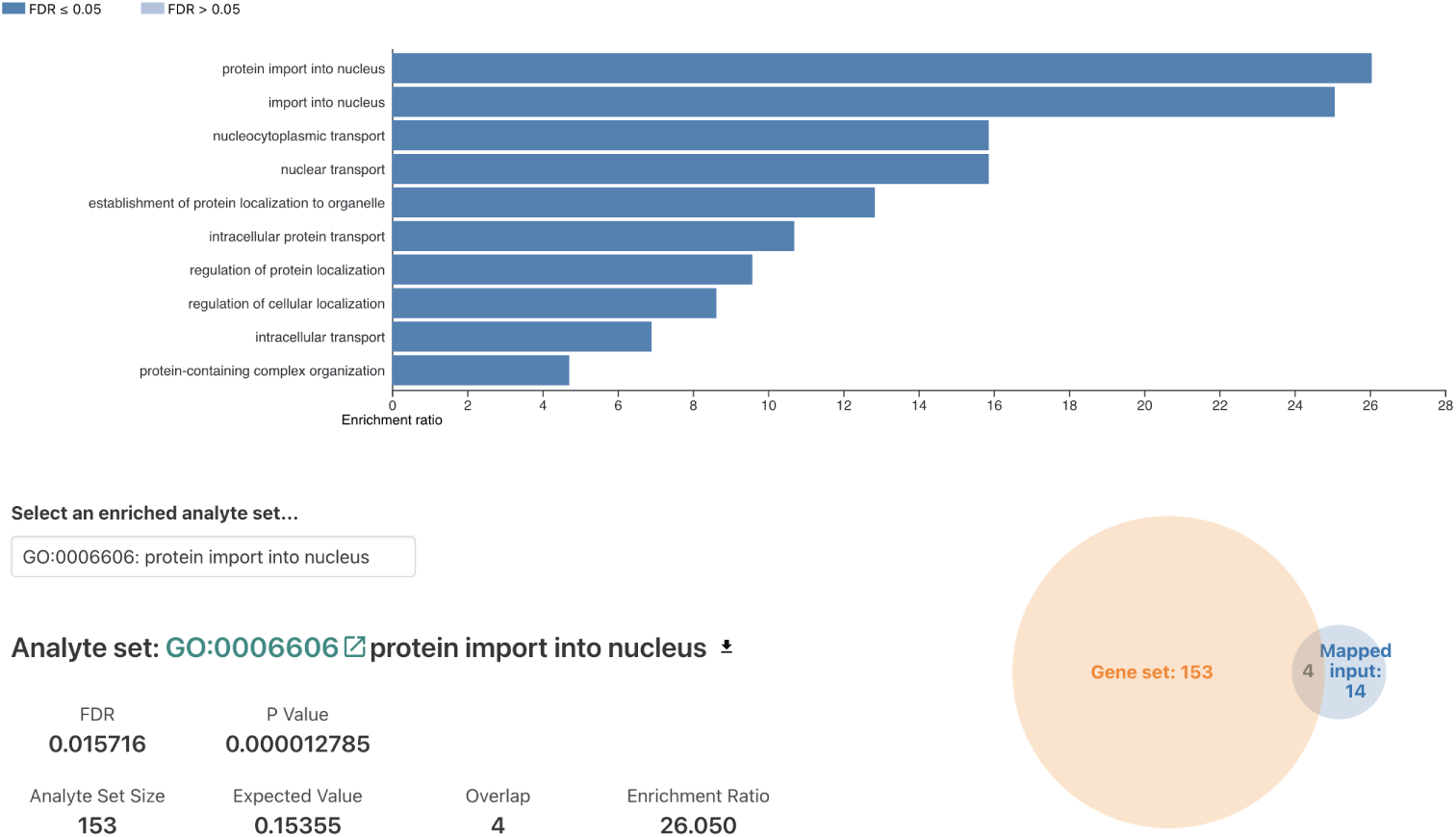
Enrichment results after performing ORA using WebGestalt for cluster 21.

**Fig. 7.**
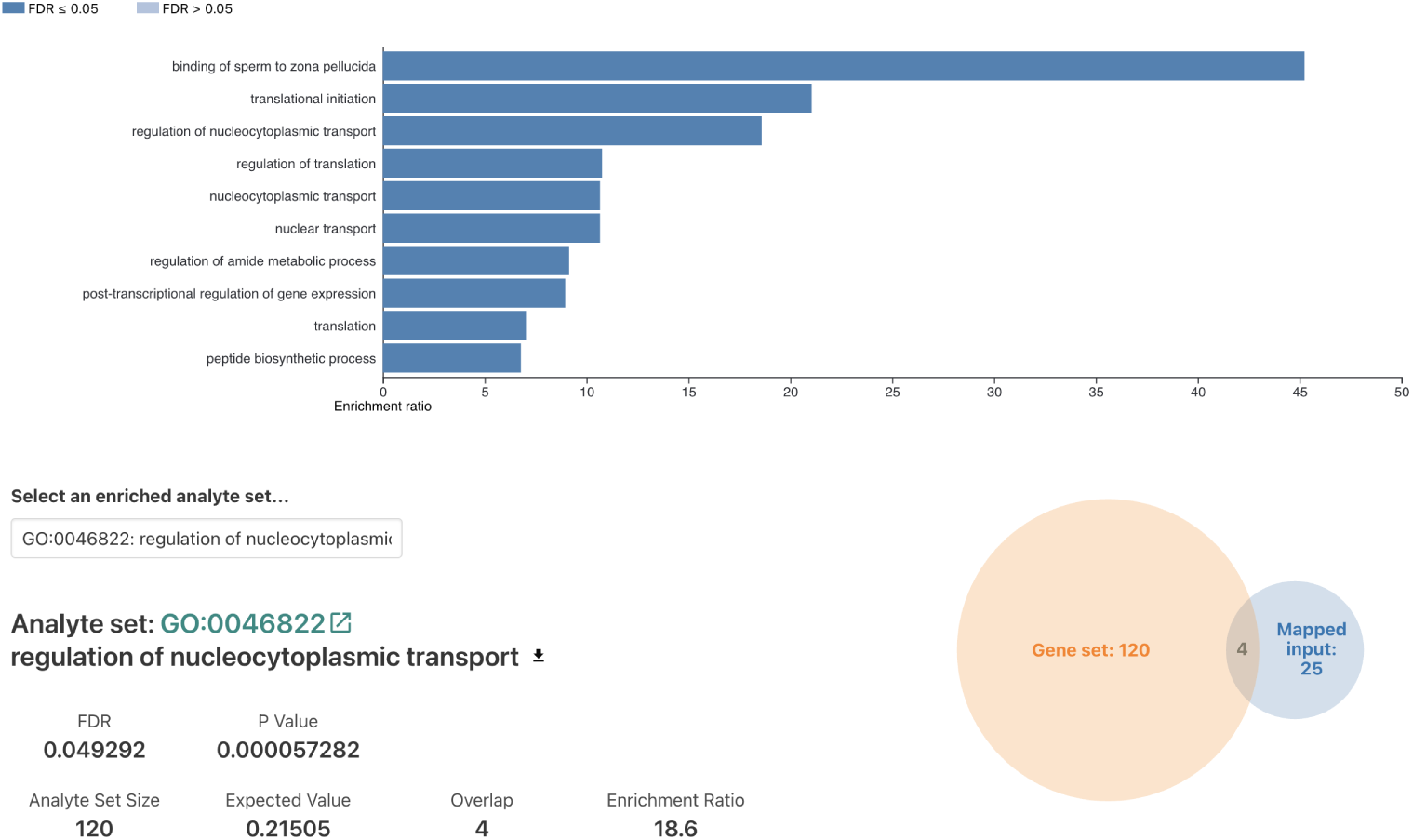
Enrichment results after performing ORA using WebGestalt for cluster 25. the semantic score values for each cluster by modulating intensity and introducing

**Fig. 8.**
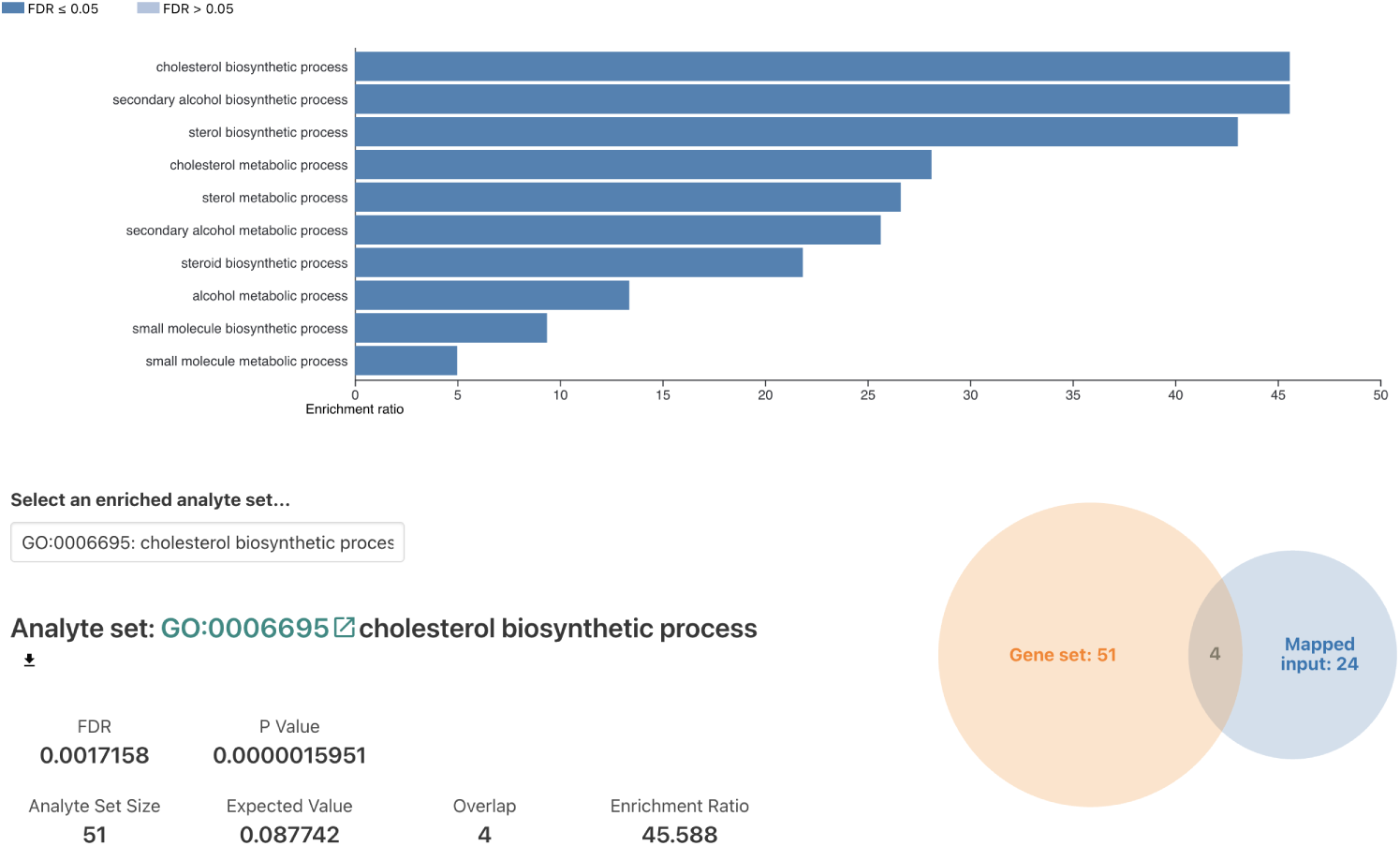
Enrichment results after performing ORA using WebGestalt for cluster 40.

Functional enrichment analysis was performed on each of the six clusters produced. However, only three clusters, cluster labels 21, 25, and 40, showed significant enrichment results (FDR *<* 0.05). The enrichment results for cluster labels 44, 49, 52 are omitted as none of the GO terms in these clusters had significant enrichment (FDR ≥ 0.05).

For semantic score calculation from the enrichment results, we implement the Wang’s method on the top three GO terms based on enrichment ratio. For eg, in cluster 21, GO:0006606 (protein import into nucleus), GO:0051170 (import into nucleus) and GO:0006913 (nucleocytoplasmic transport) are the three highly enriched GO terms. Similarly for cluster 25, GO:0007339 (binding of sperm to zona pellucida), GO:0006413 (translational initiation), GO:0046822 (regulation of nucleocytoplasmic transport) and for cluster 40, GO:0006695 (cholesterol biosynthetic process), GO:1902653 (secondary alcohol biosynthetic process), GO:0016126 (sterol biosynthetic process) are respectively the top enriched biological processes. The semantic scores obtained using Wang’s method are summarised in Table 1

**Table 1.** Semantic scores and GO terms associated with selected gene clusters.

| Cluster | Semantic Score | GO Terms |
| --- | --- | --- |
| 21 | 0.612 | GO:0006606, GO:0051170, GO:0006913 |
| 25 | 0.079 | GO:0007339, GO:0006413, GO:0046822 |
| 40 | 0.597 | GO:0006695, GO:1902653, GO:0016126 |
| 44 | 0.827 | GO:0006305, GO:0006306, GO:0006304 |
| 49 | 0.730 | GO:0051085, GO:0051084, GO:0006458 |
| 52 | 0.496 | GO:0043153, GO:0009648, GO:0009649 |

### 5.3 Generated Music

Corresponding to each of these clusters, music has been generated using the three distinct foreground music approaches – “Chords”, “Mean”, “Delayed”, mentioned earlier. Each approach translates the gene expression data into a unique auditory pattern, allowing us to interpret the temporal dynamics of gene expression through sound. In addition, we combined the background music using the semantic similarity scores.

The foreground music for each cluster effectively represents expression pattern trends by varying pitch, with highly expressed genes rendered as high-pitched notes and lowly expressed genes as low-pitched notes. The background music encodes echoes into the waveform, with higher values resulting in stronger intensity and more prominent echoes.

The music corresponding to all the clusters are available online on the SoundCloud platform: https://on.soundcloud.com/4u9lApEMTUDsbmX5Ab

### 5.4 Mel Spectrograms of the Generated Music

A Mel spectrogram is a time–frequency representation of an audio signal that maps frequencies onto the Mel scale, which approximates how humans perceive pitch. In the context of our project, Mel spectrograms are particularly useful for visually analyzing the structure and intensity of both the foreground and background layers of the musified gene expression data. Figure 9 presents the Mel spectrograms for Cluster 21, illustrating one original semantic score of 0.615 for the cluster alongside five hypothetical scores—0.05, 0.2, 0.45, 0.8, and 0.95—to demonstrate how variations in semantic similarity influence the resulting music. These spectrograms provide a visual representation of how both the foreground and background layers respond to different semantic scores. Notably, from a score of 0.05 up to around 0.5, the intensity of the foreground music—generated using the chords method—remains relatively consistent, reflecting the stable encoding of expression dynamics. In contrast, the background music gradually intensifies within this range, mirroring the increasing semantic similarity through louder and denser audio textures. However, beyond the threshold of 0.5, the pattern reverses: the foreground music begins to soften in intensity, while the background layer introduces progressively stronger echo effects. This transition highlights how the system modulates auditory characteristics to convey shifts in the biological significance of the cluster, creating a layered experience where listeners can intuitively sense changes in both expression dynamics and functional context.

**Fig. 9.**
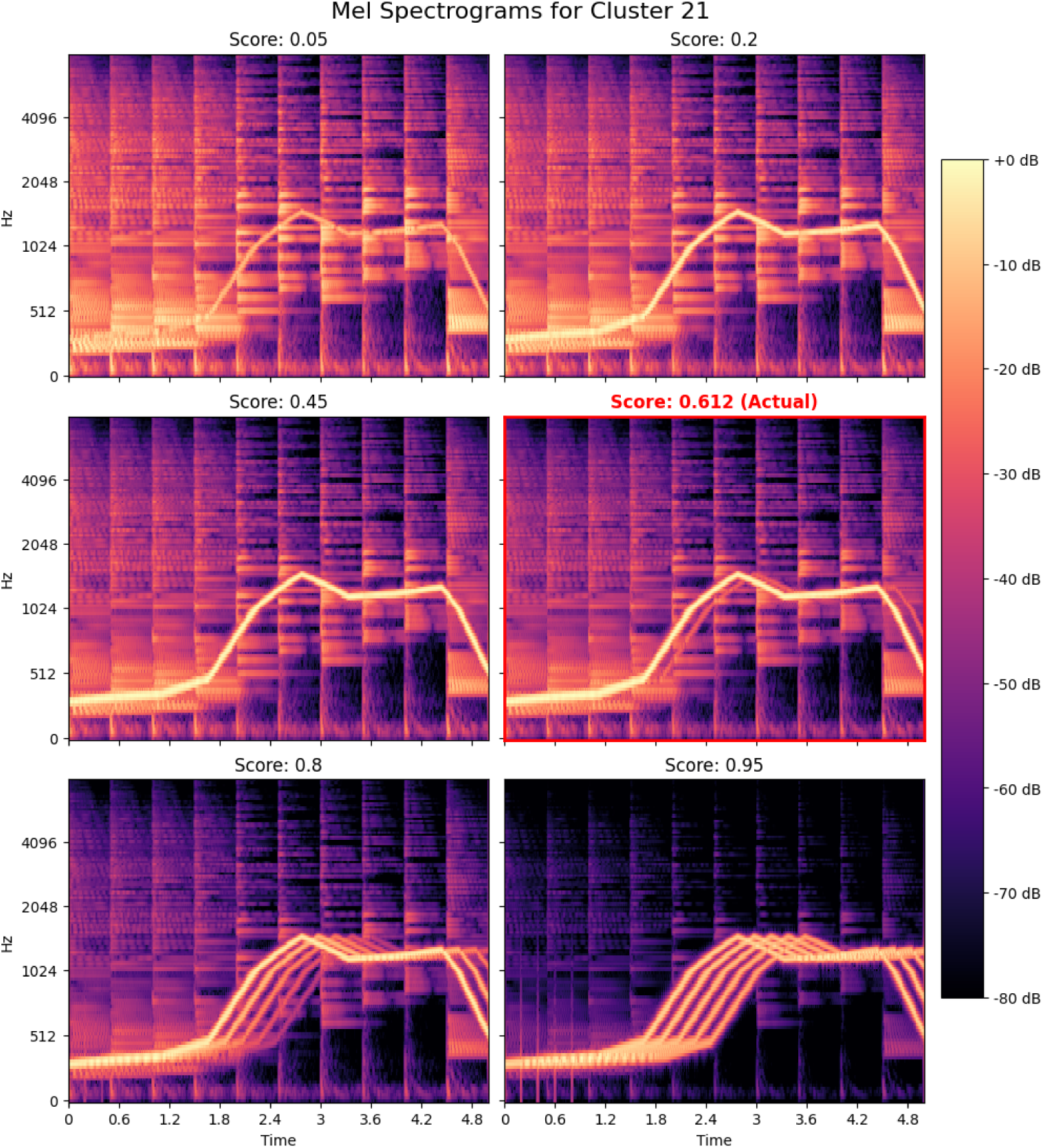
Mel spectrograms demonstrating how variations in semantic scores influence the intensity and echo characteristics of background music, as well as the foreground music dynamics, for Cluster 21. At lower semantic scores, the foreground music is more intense while the background layer remains subtle. As the semantic score increases, foreground intensity decreases, and the background music becomes more prominent, with echo effects becoming noticeable at extremely high scores.

### 5.5 User Survey Analysis

We gathered feedback from 18 individuals after playing audio files generated by our musification framework. Thirteen were computational biologists with prior experience working with gene expression data, while the remaining five were non-computational biologists, included to assess interpretability without domain knowledge. The survey comprised three sections:

- **Audio-to-Plot Mapping:** Participants were asked to listen to three audio files of clusters and match each one to the correct expression plot, evaluating how well the musification conveyed expression trends.
- **Single Gene to Cluster Identification:** Participants were provided with the audio file of a single gene from a cluster and asked to identify the correct cluster-level audio from a set of options. This was conducted separately for both the *chords* and *mean* sonification methods.
- **Echo Perception in Background Music:** Participants were presented with two smoothed audio files—one with an echo effect and one without—and were asked to compare them, assessing their ability to perceive and interpret changes in the background layer that represent semantic similarity.

The visualization in Figure 10 contains three main analyses based on survey responses. The first stacked bar chart represents participants’ ability to map audio to the correct gene expression plot. For Audio 1, 11 participants (64.7%) correctly matched the audio, while 6 (35.3%) did not. For Audio 2, 12 (66.7%) were correct and 6 (33.3%) incorrect; for Audio 3, 14 (77.8%) were correct and 4 (22.2%) incorrect. Overall, 37 out of 51 responses (72.5%) were correct. The second stacked bar chart shows the ability to identify a gene cluster based on a single gene’s audio. Using the Chords method, 12 participants (66.7%) were accurate and 6 (33.3%) were not. With the Mean method, 11 (61.1%) were accurate and 7 (38.9%) were not. Finally, the pie chart displays echo perception in background music. A significant majority, 16 out of 18 respondents (88.9%), reported that the two audios were similar but the second had an echo, while only 2 participants (11.1%) believed the audios were completely different. These results suggest that layered musification was largely interpretable and meaningful to the participants.

**Fig. 10.**
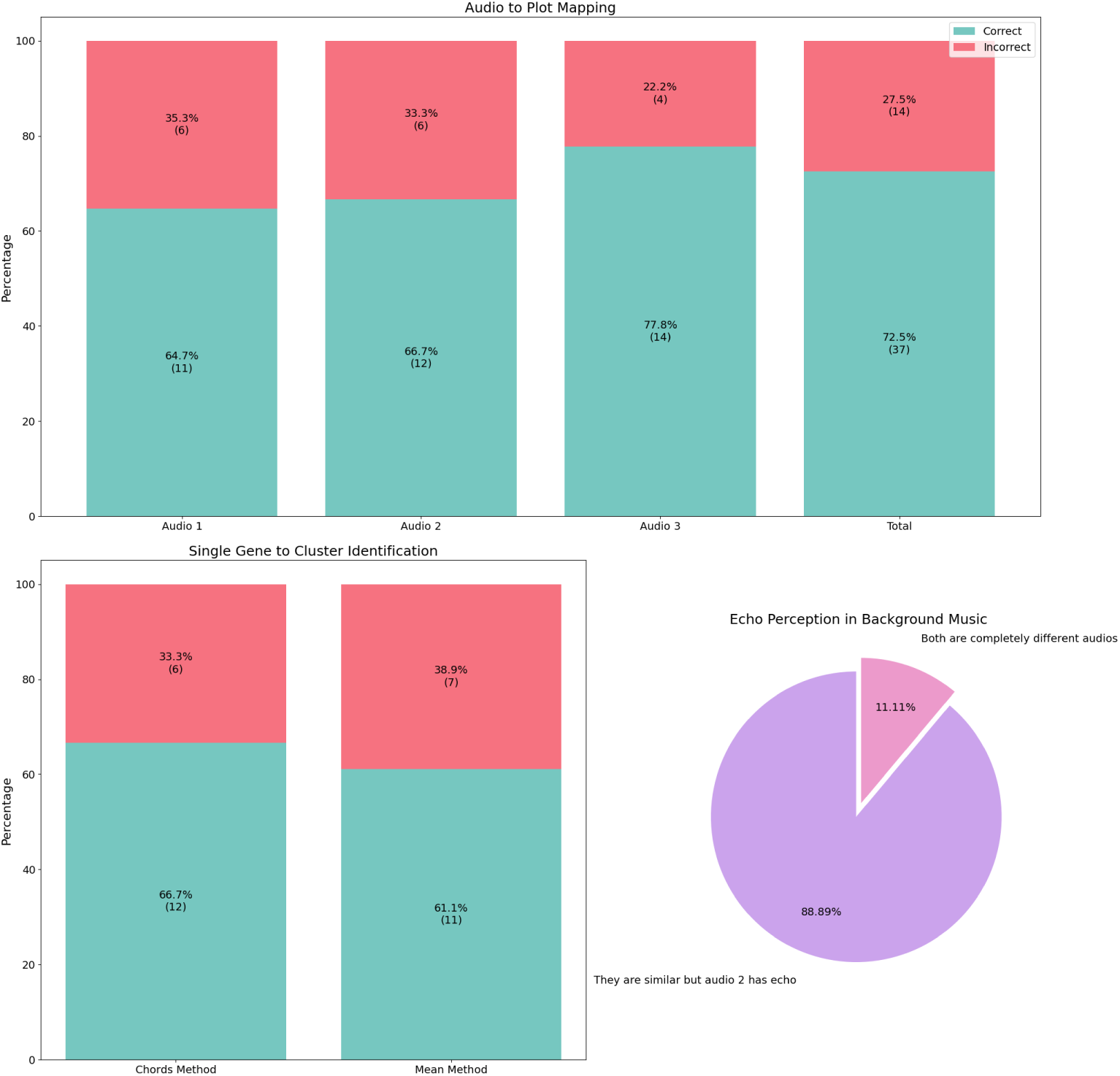
The figure summarizes participant responses from the user evaluation study. Stacked bar charts on the top indicate that a majority of users accurately mapped audio to corresponding expression plots with a total accuracy of 72.5%. Similarly, the stacked bar charts on the bottom left shows the results of correctly (green) and incorrectly (red) identified gene-to-cluster associations across two different sonification methods: Chords and Mean. The pie chart demonstrates that most participants perceived distinctions in background music using echo, validating the sensitivity of our semantic-driven audio variations. These results collectively support the interpretability and perceptual clarity of the proposed musification approach used in our study.

## 6 Conclusion

### 6.1 Summary

In this study, we explored the musification of time-series gene expression data by developing a two-layered audio representation framework. The primary layer translates temporal gene expression patterns into melody lines using multiple approaches, including chords, delayed, and means. The secondary layer incorporates biological semantics by integrating cluster-specific functional enrichment results to generate background music that reflects the biological roles of the genes. By combining musical translation with semantic depth, the method offers both auditory aesthetics and interpretability. The framework was evaluated through a structured human perception survey involving audio to graphical mapping accuracy, cluster identification, and echo audio differentiation, with 72.5%, 63.9% and 88.9% correct responses respectively, providing qualitative insights into the interpretability and relevance of musified data.

In contrast to previous works on sonification in biology—which have predominantly focused on static gene expression or protein sequence datasets—our study explores the relatively underutilized domain of time-series gene expression. Unlike studies such as M. Staege (2015) and Alterovitz et al. (2022), which translate the entire gene set into a single musical representation, often resulting in data loss and obscuring localized expression dynamics, our approach leverages gene clusters to preserve temporal patterns and expression diversity. Moreover, while earlier studies do not incorporate the biological functions of the genes in their musical interpretations, our method enriches the sonification with semantic information derived from gene ontology, thereby enhancing both biological relevance and interpretability. Additionally, our work also includes a structured, survey-based human evaluation—an important layer of validation that is notably absent in some previous gene expression sonification studies. This makes our approach distinct not only in terms of data modality and analytical depth but also in its emphasis on usability testing and user feedback.

### 6.2 Limitations and Future Work

Despite its strengths, the current work has certain limitations. First, gene expression datasets are typically large, and clustering reduces this to manageable subsets, which may result in partial biological coverage and possibly overlook some rare but important expression signals. Second, some clusters with very close semantic enrichment scores and similar expression dynamics may result in musical outputs that sound nearly identical, reducing auditory distinctiveness. This issue can be better addressed through tools like mel spectrograms, which offer detailed frequency-based visual analysis, although such methods fall outside intuitive human hearing. A third limitation involves the manual selection of parameters (e.g., tempo, instrument mapping, smoothing), which can introduce subjectivity and require further formalization for consistency and scalability.

Looking ahead, there are several promising directions for this work. Expanding the audio synthesis to incorporate multi-instrumentation and spatialized sound can significantly enhance auditory differentiation, especially for applications involving three-dimensional gene expression data, which has gained increasing attention in recent research. Incorporating machine learning to optimize cluster-to-sound mappings based on perceptual distinctiveness and biological interpretability is another potential improvement. On the research front, comparing musified outputs across different biological conditions or datasets (e.g., diseased vs. healthy) could offer novel auditory biomarkers. Further, integrating musification into educational tools and immersive visualization platforms could broaden its applicability. Lastly, extending the survey to include users with varying backgrounds—musicians, non-scientists, or individuals with impairments—can provide richer insights into usability and impact.

## 7 Code Availability

The code used for this study will be made available upon request.

## 8 Acknowledgment

The authors thank Saish Jaiswal (IIT Madras) for his varied inputs on this work. We are also grateful to the members of the BIRDS research group for their valuable feedback and support throughout the course of this study.

## References

[1] Sawe, N., Chafe, C., Treviño, J.: Using data sonification to overcome science literacy, numeracy, and visualization barriers in science communication. Frontiers in Communication 5 (2020) 10.3389/fcomm.2020.00046

[2] Fovino, L.G.N., Zanella, A., Grassi, M.: Evaluation of the effectiveness of sonification for time series data exploration (2024). https://arxiv.org/abs/2402.09953

[3] Braun, R., Tfirn, M., Ford, R.M.: Listening to life: Sonification for enhancing discovery in biological research. Biotechnology and Bioengineering 121(10), 3009–3019 (2024) 10.1002/bit.28729 https://analyticalsciencejournals.onlinelibrary.wiley.com/doi/pdf/10.1002/bit.28729

[4] Mainka, S., Schroll, A., Warmerdam, E., Gandor, F., Maetzler, W., Ebersbach, G.: The power of musification: Sensor-based music feedback improves arm swing in parkinson’s disease. Movement Disorders Clinical Practice 8(8), 1240–1247 (2021) 10.1002/mdc3.13352 https://movementdisorders.onlinelibrary.wiley.com/doi/pdf/10.1002/mdc3.13352

[5] Dasilva, E.: Art, biotechnology and the culture of peace. Early DNA music project reference (John Dunn) (2004). http://www.ejbiotechnology.info/content/vol7/issue2/full/8

[6] Dunn, J., Clark, M.A.: Life music: The sonification of proteins. Leonardo 32, 25–32 (1999)

[7] Temple, M.D.: An auditory display tool for dna sequence analysis. BMC Bioinformatics 18(1), 221 (2017) 10.1186/s12859-017-1632-x

[8] Plaisier, H., Meagher, T.R., Barker, D.: Dna sonification for public engagement in bioinformatics. BMC Research Notes 14(1), 273 (2021) 10.1186/s13104-021-05685-7

[9] Yu, C.-H., Qin, Z., Martin-Martinez, F.J., Buehler, M.J.: A self-consistent sonification method to translate amino acid sequences into musical compositions and application in protein design using artificial intelligence. ACS Nano 13(7), 7471–7482 (2019) 10.1021/acsnano.9b02180

[10] Martin, E.J., Meagher, T.R., Barker, D.: Using sound to understand protein sequence data: new sonification algorithms for protein sequences and multiple sequence alignments. BMC Bioinformatics 22(1), 456 (2021) 10.1186/s12859-021-04362-7

[11] Staege, M.S.: A short treatise concerning a musical approach for the interpretation of gene expression data. Scientific Reports 5(1), 15281 (2015) 10.1038/srep15281

[12] Staege, M.S.: Gene expression music algorithm-based characterization of the ewing sarcoma stem cell signature. Stem Cells International 2016(1), 7674824 (2016) 10.1155/2016/7674824 https://onlinelibrary.wiley.com/doi/pdf/10.1155/2016/7674824

[13] Staege, M.S., Hattenhorst, U.E., Neumann, U.E., Hutter, C., Foja, S., Burdach, S.: Dna-microarrays as tools for the identification of tumor specific gene expression profiles: applications in tumor biology, diagnosis and therapy. Klinische Padiatrie 215(3), 135–139 (2003) 10.1055/s-2003-39371. Research Support, Non-U.S. Gov’t

[14] Alterovitz, G., Yuditskaya, S.: Musical gene expression: Abstracting high-dimensional gene dynamics. J. Biotechnol. Biomater. 12, 280 (2022) 10.4172/2155-952X.1000280

[15] Huang, D.W., Sherman, B.T., Tan, Q., Collins, J.R., Alvord, W.G., Roayaei, J., Stephens, R., Baseler, M.W., Lane, H.C., Lempicki, R.A.: The david gene functional classification tool: a novel biological module-centric algorithm to functionally analyze large gene lists. Genome Biology 8(9), 183 (2007) 10.1186/gb-2007-8-9-r183. PMID: 17784955; PMCID: PMC2375021

[16] Reimand, J., Arak, T., Adler, P., Kolberg, L., Reisberg, S., Peterson, H., Vilo, J.: g:profiler—a web server for functional interpretation of gene lists (2016 update). Nucleic Acids Research 44(W1), 83–89 (2016) 10.1093/nar/gkw199. PMID: 27098042; PMCID: PMC4987867

[17] Kuleshov, M.V., Jones, M.R., Rouillard, A.D., Fernandez, N.F., Duan, Q., Wang, Z., Koplev, S., Jenkins, S.L., Jagodnik, K.M., Lachmann, A., McDermott, M.G., Monteiro, C.D., Gundersen, G.W., Ma’ayan, A.: Enrichr: a comprehensive gene set enrichment analysis web server 2016 update. Nucleic Acids Research 44(W1), 90–97 (2016) 10.1093/nar/gkw377. PMID: 27141961; PMCID: PMC4987924

[18] Liao, Y., Wang, J., Jaehnig, E.J., Shi, Z., Zhang, B.: Webgestalt 2019: gene set analysis toolkit with revamped uis and apis. Nucleic Acids Research 47(W1), 199–205 (2019) 10.1093/nar/gkz401. PMID: 31114916; PMCID: PMC6602449

[19] Zhao, B., Baloch, Z., Ma, Y., Wan, Z., Huo, Y., Li, F., Zhao, Y.: Identification of potential key genes and pathways in early-onset colorectal cancer through bioinformatics analysis. Cancer Control 26(1), 1073274819831260 (2019) 10.1177/1073274819831260. Erratum in: Cancer Control. 2019 Jan-Dec;26(1):1073274819840084. PMID: 30786729; PMCID: PMC6383095

[20] Huang, Y., Cui, G.: Hub Genes Related to the Pathogenesis and Progression of Colorectal Cancer and Adenoma by Integrative Bioinformatics Approaches. Preprint (Version 1) available at Research Square (2021). 10.21203/rs.3.rs-637295/v1. 10.21203/rs.3.rs-637295/v1

[21] Ward, J.H.: Hierarchical grouping to optimize an objective function. Journal of the American Statistical Association 58(301), 236–244 (1963) 10.1080/01621459.1963.10500845

[22] Gao, Z., Wang, H., Feng, G., Lv, H.: Exploring sonification mapping strategies for spatial auditory guidance in immersive virtual environments. ACM Trans. Appl. Percept. 19(3) (2022) 10.1145/3528171

[23] Benjamini, Y., Hochberg, Y.: Controlling the false discovery rate: A practical and powerful approach to multiple testing. Journal of the Royal Statistical Society: Series B (Methodological) 57(1), 289–300 (1995) 10.1111/j.2517-6161.1995.tb02031.x

[24] Yu, G., Li, F., Qin, Y., Bo, X., Wu, Y., Wang, S.: Gosemsim: an r package for measuring semantic similarity among go terms and gene products. Bioinformatics 26(7), 976–978 (2010) 10.1093/bioinformatics/btq064

[25] Wang, J.Z., Du, Z., Payattakool, R., Yu, P.S., Chen, C.-F.: A new method to measure the semantic similarity of go terms. Bioinformatics 23(10), 1274–1281 (2007) 10.1093/bioinformatics/btm087

[26] Almasoud, A.M., Al-Khalifa, H.S., Al-Salman, A.S.: Handling big data scalability in biological domain using parallel and distributed processing: A case of three biological semantic similarity measures. BioMed Research International 2019, 6750296 (2019) 10.1155/2019/6750296

[27] Lefol, Y., Korfage, T., Mjelle, R., Prebensen, C., Lüders, T., Müller, B., Krokan, H., Sarno, A., Alsøe, L., Lemonaid, C., Berdal, J.-E., Sætrom, P., Nilsen, H., Domanska, D.: Tisa: Timeseriesanalysis—a pipeline for the analysis of longitudinal transcriptomics data. NAR Genomics and Bioinformatics 5(1), 020 (2023) 10.1093/nargab/lqad020

[28] Dobin, A., Davis, C.A., Schlesinger, F., Drenkow, J., Zaleski, C., Jha, S., Batut, P., Chaisson, M., Gingeras, T.R.: Star: ultrafast universal rna-seq aligner. Bioinformatics 29(1), 15–21 (2013) 10.1093/bioinformatics/bts635

[29] Anders, S., Pyl, P.T., Huber, W.: Htseq–a python framework to work with high-throughput sequencing data. Bioinformatics 31(2), 166–169 (2015) 10.1093/bioinformatics/btu638

